# Pro-endometriosis macrophage release of IL-33 is key for endometriosis pain and lesion formation

**DOI:** 10.1101/2025.10.20.683510

**Authors:** Victor Fattori, Fernanda S. Rasquel-Oliveira, Soledad Ochoa, Eva Graf, Maria V. Bazzano, Thiha Aung, Mehmet Vural, Cynthia Kohl, Maria E. Solano, Matheus D. V. da Silva, Olivia K. Heintz, Stuart M. Brierley, Waldiceu A. Verri, Silke Haerteis, Joel Castro, Kate Lawrenson, Michael S. Rogers

## Abstract

Endometriosis is a painful gynecological inflammatory disease affecting up to 10% of females. When released by sensory neurons, calcitonin gene-related peptide (CGRP) shapes immunity, a process known as neuroimmune communication. We previously showed that nociceptor-derived CGRP polarizes macrophages into pro-endometriosis macrophages (PEMs) that mediates endometrial epithelial (endo-epi) cell proliferation and pain. However, the key mediators involved in this PEM-induced cell proliferation were unknown. Using unbiased approaches, we discovered that nociceptor-derived CGRP induces PEMs to produce IL-33. IL-33 binding to its receptor ST2 is key for endometriotic lesion growth and pain during endometriosis in mice as anti-IL-33 antibody treatment reduced evoked and spontaneous pain as well as lesion size. Chemical or genetic ablation of nociceptors or macrophages also resulted in lower levels of lesion IL-33, demonstrating a neuroimmune-driven mechanism for IL-33 production during endometriosis. In humans, we found that IL-33 is correlated with increased number of glands and fibrosis in lesions and that IL-33 expression in macrophages is also associated with genetic risk of endometriosis. We also provided evidence that suggests a dual role for IL-33 in endometriosis, in which, it is initially required for lesion formation and later for lesion maintenance only, and associated pain. Therefore, targeting IL-33/ST2 signaling may effectively treat endometriosis pain.

## INTRODUCTION

Endometriosis is a painful gynecological inflammatory disease that affects 10% of women (*1*) and transgender men (*2*) with annual health care and economic cost estimates ranging from $78–119 billion annually in the US alone (*3, 4*). Histologically, endometriotic lesions are distinguished by the presence of endometrial-type epithelial glands and stroma, frequently accompanied by hemosiderin-laden macrophages, and fibrosis. Additional components such as blood vessels, nerve fibers, smooth muscle, and immune cell infiltrates are also commonly observed. Clinically, patients often suffer from debilitating and chronic pelvic or abdominal pain, a hallmark feature of the disease. Although hormonal therapies and nonsteroidal anti-inflammatory drugs (NSAIDs) offer symptom relief for some individuals, they are ineffective for a substantial fraction of patients and their use is limited by adverse effects and contraindications in patients with comorbid conditions. Surgical excision remains a standard intervention; however, recurrence of both disease and pain is frequent (*5, 6*). This suboptimal effectiveness of current management strategies has been linked to increased reliance on opioids, with endometriosis patients demonstrating elevated risks of long-term opioid use, dependence, and overdose (*7*). Altogether, this highlights the need for novel therapeutic targets and medical interventions that can provide sustained benefits and reduce pain.

Nociceptors are specialized peripheral somatosensory neurons that respond to noxious and/or injurious stimuli leading to pain (*8–10*). Upon activation, the peptidergic nociceptors subpopulation releases neuropeptides such as calcitonin gene-related peptide (CGRP) that signal through the calcitonin receptor-like receptor (CALCRL) in complex with receptor activity modifying protein 1 (RAMP1). The release of these neuropeptides by nociceptors shapes immune cell function in the innervated tissue in a context-dependent manner. This process is known as neuroimmune communication (*8–10*).

We have previously demonstrated a role for neuroimmune communication during endometriosis (*11*). We found a nociceptor to macrophage communication via CGRP/RAMP1 signaling that polarizes macrophages to a pro-endometriosis phenotype, resulting in pro-endometriosis macrophages (PEMs). *In vitro*, we found that CGRP-induced release of mediators by PEMs leads to endometrial epithelial (endo-epi) cell growth and drives lesion formation and pain (*11*). However, the key mediators involved in this PEM-induced endometrial cell growth were unknown.

Interleukin-33 (IL-33) is a member of the IL-1 family and the sole agonist of the previous orphan receptor ST2 (*12*). Both IL-33 and ST2 are expressed by virtually all immune cells (*13, 14*). Seminal studies demonstrated that IL-33 induces pain and exacerbates disease burden, either by directly activating nociceptors or by potentiating the release of inflammatory stimulus (*15–23*). Specifically for endometriosis, genome-wide association studies (GWAS) show that variants proximal to IL-33 are associated with endometriosis risk (*24, 25*). In mice, previous work has demonstrated that IL-33 contributes to the development of endometriosis lesions (*26, 27*). But those studies used exogenous administration of IL-33 at high doses and/or surgery to induce endometriosis. Moreover, the key molecules involved in regulating IL-33 production were also not addressed. Surgery in the abdominal cavity changes the phenotype of peritoneal cells, confounding analysis of the pathophysiology of endometriosis (*28*). Additionally, exogenous administration of IL-33 at doses as low as 0.4 µg induces eosinophilia and production of Th2 cytokines (*29*), which again might be confound understanding the role of a cytokine in a disease model.

Here, using unbiased approaches such as scRNAseq and bulk RNAseq, we discovered that nociceptor-derived CGRP induces the production of IL-33 by PEMs that is key for lesion growth and pain during endometriosis in mice. In agreement, our single cell disease relevance score (scDRS) that integrates human scRNAseq annotations with GWAS (*30–32*), identified IL-33 expression in macrophages as associated with endometriosis risk. We also provide evidence that suggests a dual role for IL-33 in endometriosis, in which, it is initially required for lesion formation and later for lesion maintenance only. Furthermore, we show evidence suggesting that IL-33 mediates endometriosis-associated pain, via a mechanism involving a direct and potent activation of ST2^+^ nociceptors. Therefore, targeting IL-33/ST2 signaling might be an effective way to treat endometriosis and associated pain.

## RESULTS

### IL-33 production by PEMs promotes endometrial epithelial (endo-epi) cell proliferation

We previously showed that macrophages respond to CGRP and stimulate endo-epi cell growth (*11*). We repeated these experiments and confirmed that CGRP-induced PEMs increase the growth of endo-epi cells in a co-culture system (Fig 1A). This effect was RAMP1-dependent as treatment with rimegepant, a RAMP1 antagonist, reduced PEM-induced endo-epi cell growth (Fig 1). This demonstrates that PEMs release factors that support endometrial cell growth. To identify mediators involved in this cell growth, we performed bulk RNAseq of vehicle- or CGRP-stimulated macrophages as well as naïve endo-epi cells. Our bulk RNAseq data revealed 111 differentiated expressed genes (DEGs) in PEMs, including changes in *Il33* transcripts (Fig 1B). Using CellPhoneDB to identify which transcripts might be involved in cell-cell signaling. We examined the endo-epi cell RNAseq data to assess the mRNA expression level (in transcripts per million, TPM) for each of the receptors for differentiated expressed genes. We found that of these receptors, the IL-33 receptor ST2 (*Il1rl1*)/IL-1Rap (*Il1rap*) was highly expressed by endo-epi cells (Fig 1C, right side of table). Interestingly, ST2 was one of the few receptors expressed by endo-epi cells whose ligand was also increased in CGRP-mediated PEMs (Fig 1B), suggesting that PEM-induced endo-epi cell growth was via IL-33/ST2 signaling (Fig 1C). To confirm this, we performed a cell proliferation assay using vehicle-control- or IL-33-stimulated endo-epi cells. We found that mouse recombinant IL-33 induced endo-epi cell proliferation (Fig 1D, left graph). This increased proliferation is ST2-dependent, as treatment with an anti-ST2 antibody impaired IL-33-induced cell proliferation, in a concentration-dependent manner, vs. IgG control (Fig 1D, right graph). To understand the mechanisms involved in this cell growth, we performed bulk RNAseq of vehicle- or IL-33-stimulated endo-epi cells. Our bulk RNAseq analysis revealed a total of 408 DEGs (Fig 1E and S1). Among the upregulated genes, we found genes related to cell proliferation such as cellular communication network factor 4 (*Ccn4*), follistatin (*Fst*), and insulin-like growth factor-binding protein 7 (*Igfbp7*); while downregulated genes such as dual specificity phosphatase 1 and 22 (*Dusp1* and *Dusp22*) are involved in controlling apoptosis (Fig 1E). Finally, we confirmed that in mouse endometriosis lesions, ST2^+^ cells were also present both in the epithelium and stroma with increased presence in the epithelium (Fig 1F and G, left graph). Interestingly, we found that ST2^+^ epithelial cells were also proliferating as observed by co-localization with Ki-67^+^ cells (Fig 1F and G, right graph). Altogether, we found that IL-33/ST2 signaling is involved in cell proliferation both *in vitro* and *in vivo*.

**Figure 1.**
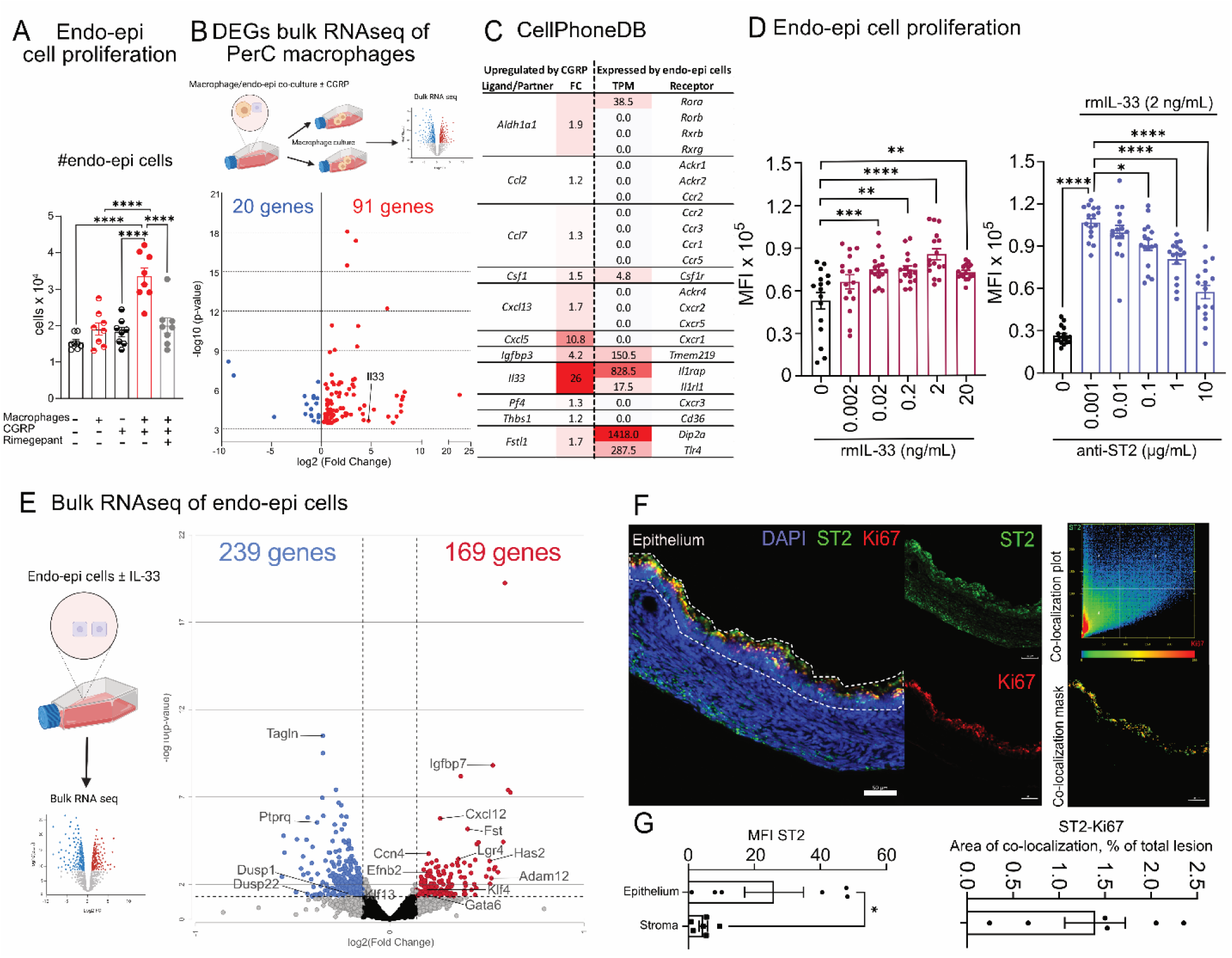
IL-33 production by PEMs promotes endometrial epithelial (endo-epi) cell proliferation. **(A)** The effect of 100 nM CGRP-treated macrophages on endo-epi cell proliferation over 24h, measured using CyQuant. Treatment with rimegepant (100 nM) or vehicle was performed 30 min. before stimulus with CGRP. Results are expressed as mean ± SEM, n = 8 (one-way ANOVA followed by Tukey’s post hoc, ****p<0.0001). **(B)** DEGs and bulk RNA-seq of peritoneal cavity macrophages stimulated with conditioned media (CM) from the vehicle- or CGRP-stimulated co-cultured cells (n = 2 for each condition). **(C)** Table displaying the fold-change (FC) of ligands or ligane-generating enzymes that were significantly upregulated upon CGRP treatment of macrophages (left) and the expression of cognate receptors expressed by endo-epi cells (right) in transcripts per million (TPM). **(D)** The effect of rmIL-33 (left bar graph) and anti-ST2 antibody (right bar graph) on endo-epi cell proliferation over 24h, measured using CyQuant. Results are expressed as mean ± SEM, n = 16 (one-way ANOVA followed by Tukey’s post hoc, ****p<0.0001; ***p<0.001; **p<0.01; *p<0.05). MFI: mean fluorescence intensity;. **(E)** DEGs from bulk RNA-seq of vehicle- or IL-33-stimulated endo-epi cells (n = 2 for each condition). **(F)** Immunofluorescence staining of endometriosis lesions displaying ST2 expression (green staining, top inset) and Ki-67 (red staining, bottom, inset). Far right insets display co-localization plot (top inset) and co-localization mask (bottom inset). **(G)** Quantification of ST2 staining (left bar graph) and area of co-localization between ST2 and Ki-67 (right bar graph). Scale bar 50µm.

### IL-33 expression in macrophages is associated with endometriosis risk in patients

Having observed a crucial role for IL-33/ST2 signaling in mouse lesions, we next wanted to determine whether IL-33 was also in human lesions. To that end, we examined 13 lesions from patients with a mean age of 34 years old (Supplementary Table 1). We observed widespread immunohistochemical staining for IL-33 throughout the lesions, particularly in both the nucleus and cytoplasm of the epithelium of endometrial glands, in the stroma of fibrotic regions, and the endothelium of blood vessels (Fig 2A). We also found Ki67^+^ epithelial cells in human lesions (Fig 2A), though to a lesser extent than in mouse lesions. Nevertheless, this indicates that our mouse endometriosis lesions recapitulate features observed in human lesions (Fig 1F). Importantly, overall IL-33 staining is positively associated with the number of glands in lesions (Fig 2B, top left graph). As fibrosis is an important hallmark of endometriosis lesions (*33*) and IL-33 is known to drive fibrosis in different organs (*34*), we next evaluated the relationship between IL-33 and fibrosis. Within each biopsy, the relative extension of fibrosis, characterized by typical dense clusters of elongated cells in the vicinity of the endometrium-like lesion, was scored on a scale from 0 to 4. We found that IL-33 staining in glands is positively associated with its staining in the fibrotic areas (Fig 2B, top right graph). We also found that the IL-33 is positively associated with overall fibrotic score (Fig 2B, bottom left graph), further suggesting that IL-33 can drive fibrosis. The extent of fibrosis demonstrated a trend toward a positive correlation with the duration since symptom onset (ρ = 0.671, p = 0.06, n = 8). Although this result did not reach statistical significance, it suggests that longer symptom duration may be associated with greater fibrotic development, supporting the hypothesis of a progressive fibrotic process in ectopic lesions. We also evaluated correlation with and among other patient demographic, symptom, and histological factors, but found no significant correlations (Supplementary Table 2). Altogether, this indicates that IL-33 is correlated with a higher number of glands as well as a greater extent of fibrosis in human lesions.

**Figure 2.**
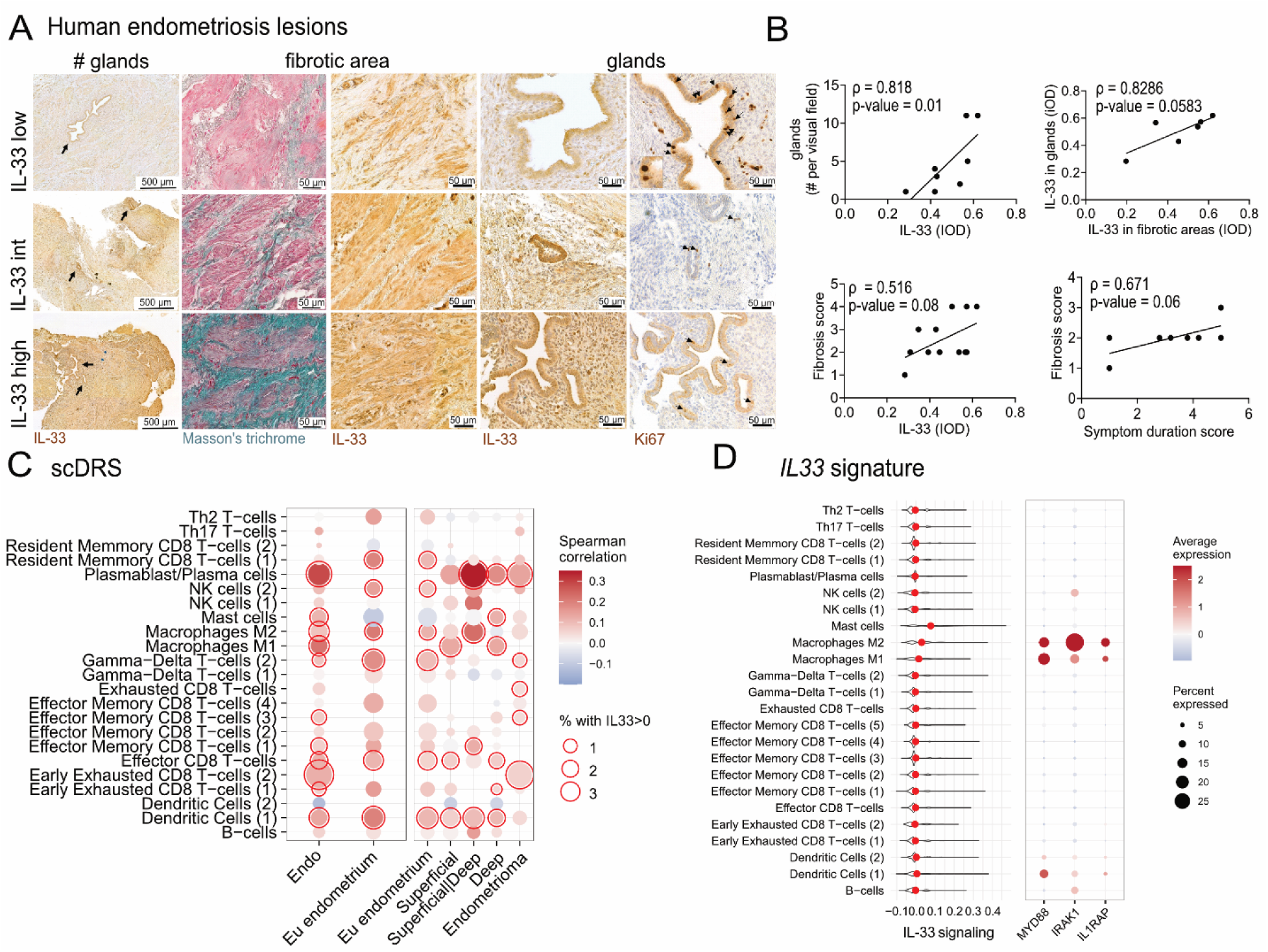
Patient IL-33 expression is associated with lesion glandularity, fibrosis, and endometriosis risk. **(A)** Immunohistochemistry (IHC) staining for IL-33 in human lesions and Masson’s trichrome staining. Images show the presence of IL-33^+^ glands (black arrows) and IL-33 within fibrotic regions and around endometrial glands. **(B)** Correlation between number of glands and IL-33 staining (top left graph), IL-33 staining in glands and IL-33 staining in fibrotic areas (top right graph), fibrosis score and IL-33 staining (bottom left graph), and fibrosis score and symptom duration (scored as: 1 = 0–6 months, 2 = 6–12 months, 3 = 1–3 years, 4 = 3–10 years, 5 = >10 years). Results are presented as Spearman’s correlation coefficients (ρ), statistical significance was set at *p < 0.05 (two-tailed). IOD = integrated optical density. **(C)** scDRS for IL-33 in immune cells. Left table displays the expression of IL-33 in the immune cell clusters with disease risk (aggregating lesion subtypes as “endo”). Red border highlights significant raw score for IL-33 and the presence of endometriosis lesions (endo). Right table displays the expression of IL-33 in the immune cell clusters with endometriosis lesion subtype risk. Red border highlights significant raw score for IL-33 and the different types of endometriosis lesions and eutopic endometrium. Correlation was not calculated for clusters below 50 cells and p<0.05 was considered significant. **(D)** Transcriptional signature of IL-33-mediated signaling per cell cluster. Red dots mark the mean of the distribution. Right table displays selected IL-33/ST2 downstream signaling pathways markers.

GWAS show that IL-33 is associated with the risk of developing endometriosis (*24, 25*). To expand such findings, we then used the single cell disease relevance score (scDRS) to specifically pinpoint the cell type associated with such risk. scDRS is an algorithm that integrates single cell expression data with germline risk associations from large genome-wide association studies (*35*). Here we integrated GWAS data from a meta-analysis of 450,668 controls and 23,492 endometriosis cases with single cell data from endometrium and endometriosis lesions from a cohort of 21 patients (*30, 32, 36*). Focusing on immune cells, we found that IL-33 expression in several cells such as plasma cells, dendritic cells, effector CD8 T-cells, mast cells, and “M1” and “M2” macrophages is significantly correlated with endometriosis risk score (Fig 2C, left graph). We then sought to determine whether IL-33 expression in these immune cells would be particularly associated with the risk of developing different types of endometriosis lesions (e.g., superficial vs. deep vs. endometrioma, etc.). We found that, except for endometriomas (ovarian lesions), IL-33 signal in macrophages was widespread for different types of endometriosis lesions, indicating that it might be an important driver of lesion formation or maintenance (Fig 2C, right graph). IL-33 signal in other cell types such as plasma cells and dendritic cells showed a similar pattern, indicating these cells could also play a role in IL-33-driven lesion formation. To further expand these results, we next evaluated a transcriptional signature of IL-33 signaling (GO:0038172, see Supplementary Material). We found that IL-33-mediated signaling as well as some of the downstream markers of IL-33/ST2 signaling were enriched in macrophages (Fig 2D), further indicating that IL-33 in macrophages could play a role in endometriosis lesion formation or maintenance.

### TRPV1^+^ nociceptors control IL-33 production by macrophages

We previously demonstrated that TRPV1^+^ nociceptors control monocyte recruitment during endometriosis, thereby facilitating lesion growth and pain (*11*). However, whether these lesion-recruited macrophages produce IL-33 and are sensitive to nociceptor ablation was unknown. Therefore, we bred *Trpv1*-cre mice with Rosa26 *lox*-STOP-*lox* DTA mice (ROSA-DTA) to create *Trpv1*cre*Dta* nociceptor-ablated mice and Cre^−^ littermate control (LM) mice. Using immunofluorescence of mouse lesions, we found that nociceptor-ablated mice had fewer F4/80^+^ macrophages as well as less IL-33 (Fig 3A). In agreement, we found a reduction in the pixel count as well as in the co-localization area between F4/80 and IL-33 upon ablation of TRPV1^+^ nociceptor (Fig 3B). This indicates that TRPV1^+^ nociceptors control the presence of F4/80^+^IL-33^+^ macrophages in the lesion.

**Figure 3.**
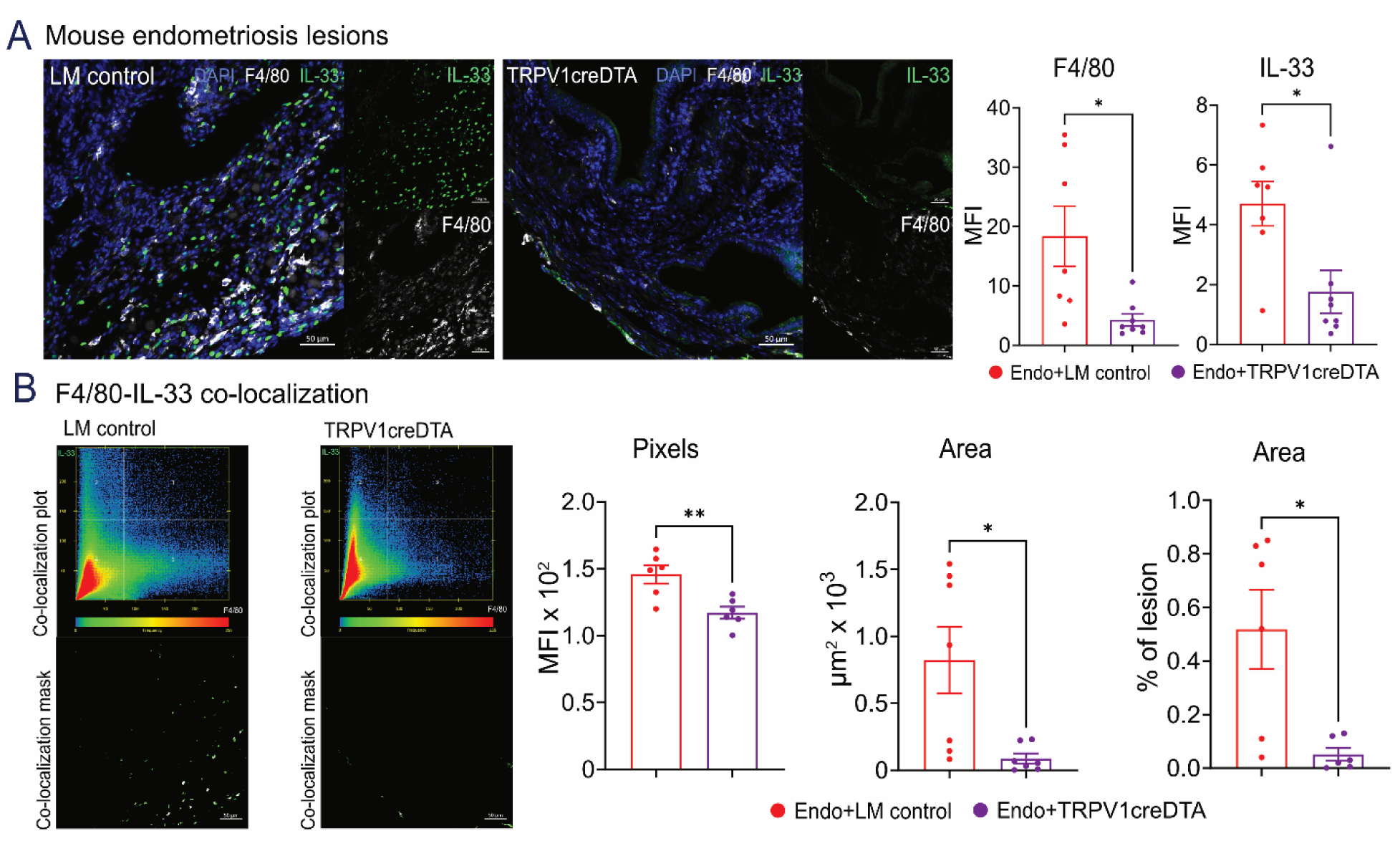
TRPV1^+^ nociceptors control IL-33 production by macrophages. **(A)** Immunofluorescence staining of endometriosis lesions from LM control and *Trpv1*cre*Dta* mice displaying IL-33 (green staining, top inset) and F4/80^+^ macrophages expression (white staining, bottom inset). Left bar graph displays quantification of mean fluorescence intensity (MFI) for F4/80 and right bar graph for IL-33. **(B)** Co-localization plot (top inset) and co-localization mask (bottom inset) of LM control and *Trpv1*cre*Dta* mice. Left bar graph displays pixel count of F4/80-IL-33 co-localization, while middle and right bar graphs display area of co-localization in µm^2^ and percentage of lesion, respectively. Scale bar 50µm. Results are expressed as mean ± SEM, n = 5 (Student’s t-test, **p<0.01; *p<0.05).

### Nociceptor to macrophage communication via CGRP/RAMP1 signaling drives IL-33 production

We have previously observed that endometriosis lesions are supported by TRPV1^+^ nociceptors which release CGRP, which then acts on local macrophages (*11*). Our next step was to determine the extent to which IL-33 production was driven by this nociceptor to macrophage communication. For that, we used chemical and genetic approaches to ablate nociceptors (Fig 4A) or macrophages (Fig 4B) as well as four different FDA-approved drugs that target either CGRP or RAMP1 (Fig 4C) to measure IL-33 levels in the lesions. We first performed chemical ablation of nociceptors using subcutaneous injection of resiniferatoxin (RTX), an agonist for TRPV1 (*39, 40*) (Fig 4A, left graph). Upon chemical ablation of TRPV1^+^ nociceptors, we found a reduction in IL-33 levels in the lesion when compared to vehicle-treated mice (Fig 4A, left graph). In agreement, genetic ablation of the same population using *Trpv1*cre*Dta* mice reduced IL-33 levels as well (Fig 4A, right graph), further indicating that nociceptors regulate IL-33 production.

**Figure 4.**
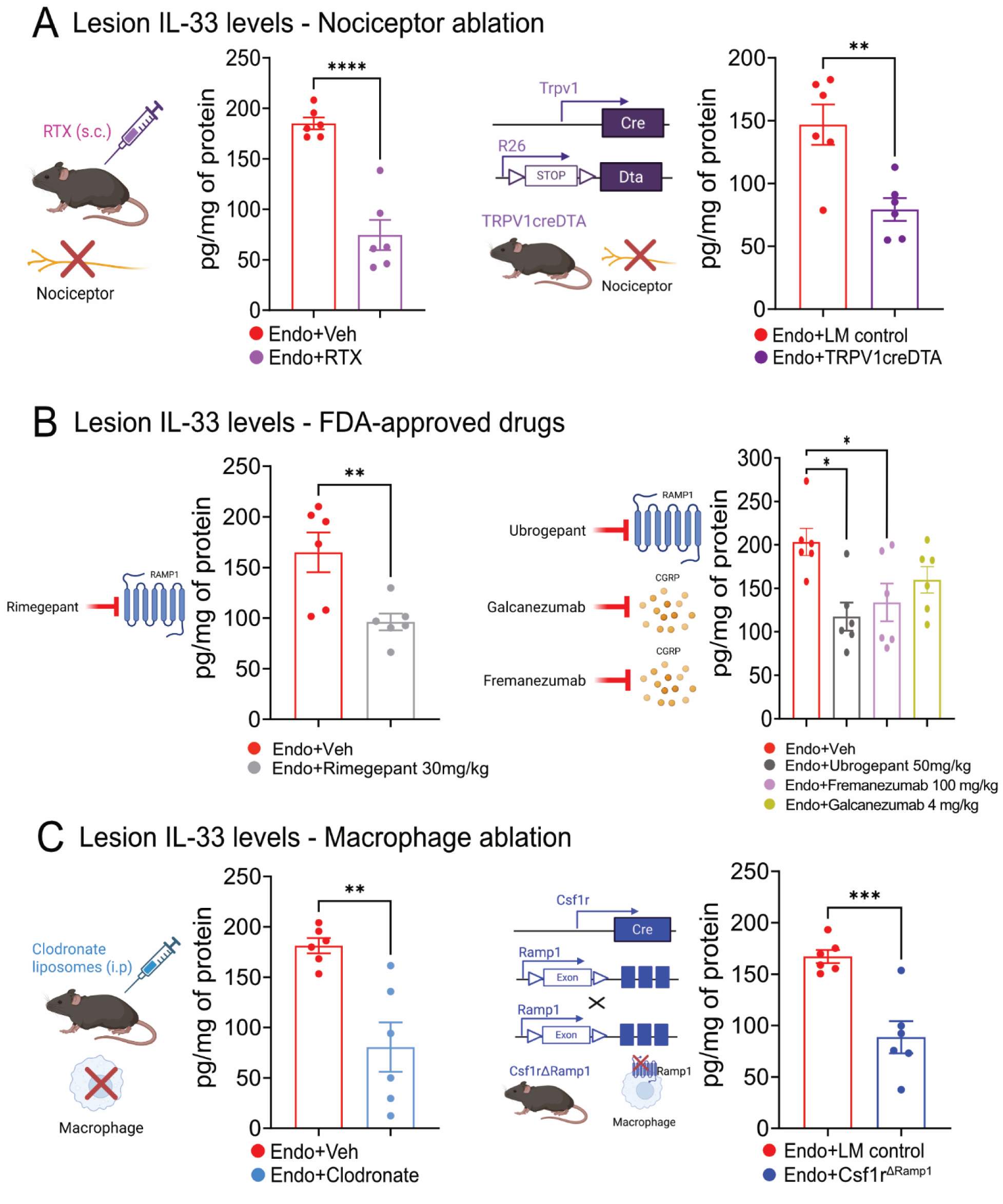
Nociceptor to macrophage communication via CGRP/RAMP1 signaling drives lesion IL-33 production. **(A)** ELISA for IL-33 in lesions of nociceptor-ablated mice. Schematic displays chemical ablation of nociceptors using RTX (left) and animal breeding to generate *Trpv1*cre*Dta* mice (right). **(B)** ELISA for IL-33 in lesions of mice treated with FDA-approved drugs. Schematics display drug targets; rimegepant (left) and ubrogepant (right) are small molecules that target RAMP1, while fremanezumab and galcanezumab (right) are monoclonal antibodies against CGRP. Left bar graph displays measurements in vehicle- and rimegepant (30 mg/kg)-treated mice at 42 dpi. Right bar graph displays measurements in vehicle- and ubrogepant (50 mg/kg)-, fremanezumab (100 mg/kg)-, or galcanezumab (4 mg/kg)-treated mice at 56 dpi. **(C)** ELISA for IL-33 in lesions of macrophage-ablated mice. Schematic displays chemical ablation of macrophages using clodronate liposomes (left) and animal breeding to generate Csf1r^Δ*Ramp1*^ mice (right). Results are expressed as mean ± SEM, n = 5 (one-way ANOVA followed by Tukey’s post hoc, ****p< 0.0001; ***p<0.001; **p<0.01; *p<0.05).

We next used FDA-approved drugs that target either CGRP (galcanezumab and fremanezumab, monoclonal antibodies) or RAMP1 (rimegepant or ubrogepant, small molecules). We found these drugs also reduced IL-33 levels (Fig 4B), supporting the notion that CGRP/RAMP1 signaling drives IL-33 production.

Then, we chemically ablated peritoneal cavity macrophages using clodronate liposomes (Fig 4B, left graph). Peritoneal-cavity-macrophage-depleted mice displayed lower levels of IL-33 when compared to vehicle-treated mice (Fig 4C, left graph). Finally, to confirm that this reduction in IL-33 was a result of nociceptor CGRP to RAMP1 signaling in macrophages, we bred a macrophage-specific cre line (*Csf1r*-cre) with *Ramp1^fl/fl^*to delete *Ramp1* from macrophages (Fig 4C, right graph Csf1r^ΔRamp1^). Mice with *Ramp1*-deleted macrophages showed reduced IL-33 production when compared to LM controls, demonstrating that nociceptor to macrophage communication via CGRP/RAMP1 signaling drives IL-33 production (Fig 4C, right graph).

### IL-33/ST2 signaling is required for lesion establishment and growth

Having observed that IL-33 induces endo-epi cell proliferation (Fig 1D) and that IL-33 staining correlates with the number of glands in human lesions (Fig 3B, left graph), we next sought to determine whether IL-33 signaling is required for lesion formation. We first determined the dynamics of IL-33 production in the peritoneal cavity wash upon endometriosis induction by ELISA. We found that IL-33 peaks 3 days post induction and then returns to baseline (Fig 5A, left bar graph), indicating that in the peritoneal cavity a burst of IL-33 is followed by a normalization of its levels during endometriosis induction. Interestingly, we detected IL-33 levels in the lesions at time points in which peritoneal cavity IL-33 levels were at baseline, e.g. 14 and 28dpi (Fig. 5A, right bar graph). This indicates that IL-33 may be a crucial lesion regulator. To confirm that IL-33/ST2 signaling is required for lesion formation, we induced endometriosis using uterine horns of WT or mice lacking *St2* (Fig 5B). When endometriosis was induced with uterine horns of *St2*-KO mice, we observed a reduction in the number of lesions, lesion size, and burden (Fig 5C). We also found that remaining lesions had less IL-33 (Fig 5D). To expand these results, we next used a pre-treatment schedule (i.e., treatment before endometriosis induction) with an anti-IL-33 antibody (Fig 5E). We found that anti-IL-33 antibody treatment reduced evoked pain as observed by an increase in the mechanical threshold (Fig 5F) and also reduced both abdominal contortions and squashing (Fig 5G). Pre-treatment with anti-IL-33 antibody also reduced the number of lesions, lesion size and burden (Fig 5H). This indicates that IL-33/ST2 signaling is required for lesion formation, establishment, and endometriosis-associated pain.

**Figure 5.**
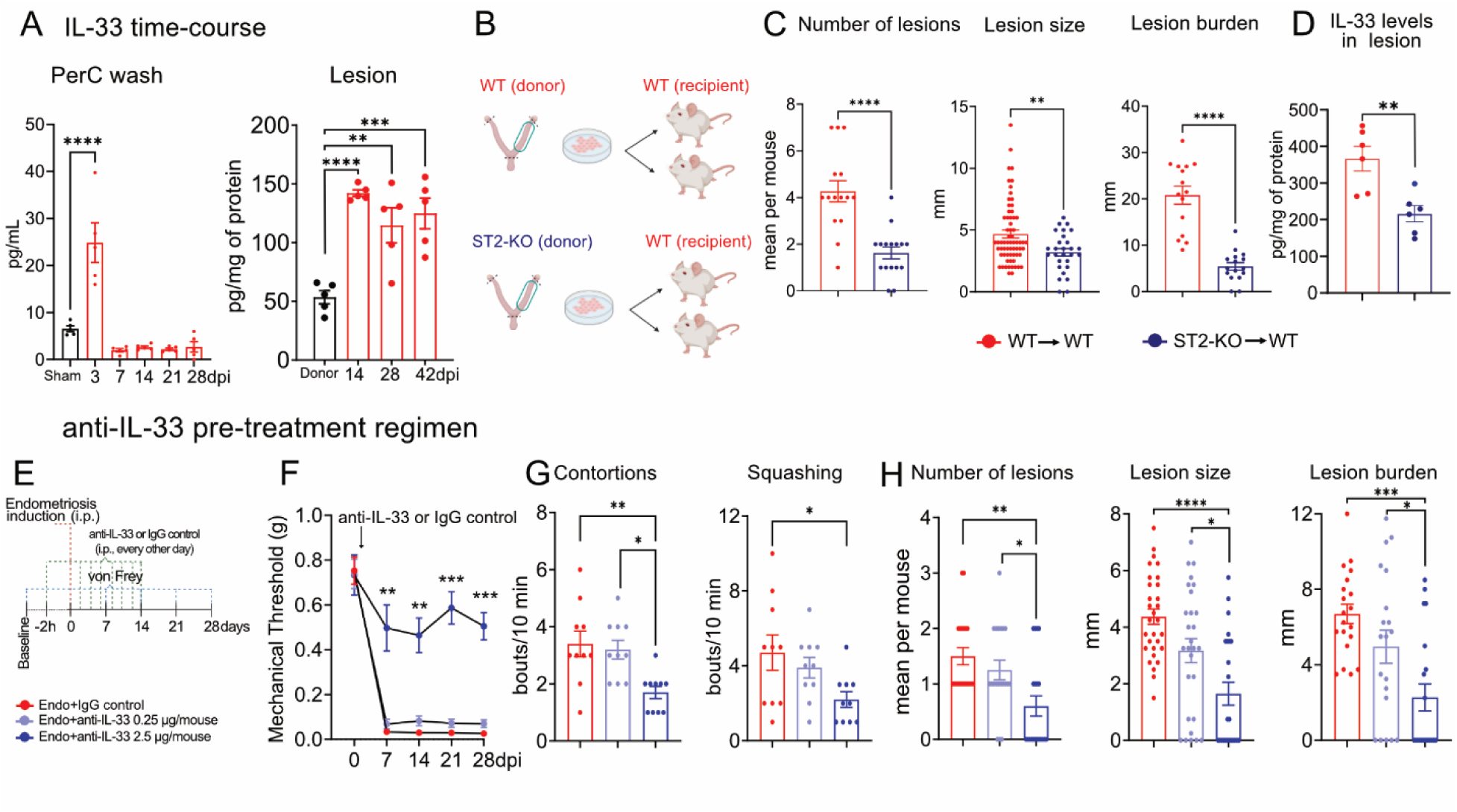
IL-33/ST2 signaling is required for lesion establishment and generation of endometriosis pain. **(A)** Time-course of IL-33 content (ELISA) in the peritoneal cavity wash (left bar graph) and lesions (right bar graph). Results are expressed as mean ± SEM, n = 5 (one-way ANOVA followed by Tukey’s post hoc, ****p< 0.0001; ***p<0.001; **p<0.01). **(B)** Schematic for the experiments involving the use of *St2*-KO mice as tissue donors. **(C)** Number of visible lesions was calculated as sum of total lesions per mouse while lesion size was measured in the lesions as the average of perpendicular diameters. Lesion burden was calculated as the sum of all lesion size in a given mouse. **(D)** ELISA for IL-33 in the lesions of mice with endometriosis induced using uterine horns of WT or *St2*-KO as tissue donors. Results are expressed as mean ± SEM, n = 17 (Student’s t-test, ****p< 0.0001; **p<0.01). **(E)** Schematic displaying the pre-treatment protocol for the anti-IL-33 antibody. **(F)** Abdominal mechanical hyperalgesia measured before lesion induction (zero) and weekly thereafter using von Frey filaments. Results are presented as mean ± SEM of mechanical threshold, n = 10 mice per group (two-way repeated-measures ANOVA followed by Tukey’s post hoc, ***p<0.001; **p<0.01). **(G)** Spontaneous pain measured by abdominal squashing and abdominal contortion. **(H)** Number of visible lesions, lesion size, and lesion burden as described above. Results are expressed as mean ± SEM, n = 10 (one-way ANOVA followed by Tukey’s post hoc, ****p< 0.0001; **p<0.01; *p<0.05).

### IL-33/ST2 signaling drives endometriosis-associated pain

We next determined whether anti-IL-33 therapy could effectively intervene in ongoing disease (e.g., treatment starting at 29 dpi, when lesions are fully developed in our mouse model, Fig 6A). Like the pre-treatment protocol, we found that post-treatment with anti-IL-33 antibody also reduced evoked pain as evidenced by an increase in the mechanical threshold (Fig 6B) and decreases in both abdominal contortions and squashing (Fig 6C). However, with this post-treatment protocol, we did not observe significant changes in lesion development (Fig 6D). Previous works have demonstrated that nociceptors express ST2 (*20, 22, 34, 41*) and that injection of recombinant IL-33 induces pain (*16–18, 42, 43*), indicating that IL-33 directly activates nociceptors. Based on that data from the literature and our own data, IL-33 might have a dual role in endometriosis. At early time points, IL-33 participates in lesion establishment and once lesions are formed, IL-33 increases endometriosis pain. To address the question of whether IL-33 can mediate pain during endometriosis through direct activation of nerve fibers, we first performed immunofluorescence of mouse endometriosis lesions. We found that they contain ST2^+^ nerve fibers as observed by double staining of ST2 with PGP9.5 (pan neuronal marker, Fig 6E). We then performed calcium imaging of DRG neurons from mice with and without endometriosis, to investigate the effect of IL-33 on sensory neuron’s activity. For that, we used NaV1.8-GCaMP6s mice which express the calcium sensor GCaMP6s in NaV1.8^+^ nociceptors (Fig 6F), as previously described (*44*). We found that IL-33 directly activates a proportion of DRG neurons cultured from both sham and endometriosis-lesion bearing mice (Fig 6G and H). Specifically, we found that endometriosis induced a >4-fold increase in the number of IL-33-responsive nociceptors with 122 out of 1006 responding DRG neurons in endometriosis mice vs 28 out of 912 responding DRG neurons in sham mice (Fig 6I). We also found that the overall magnitude of the response to IL-33 was higher in DRG neurons from mice with endometriosis, compared to those cultured from sham mice (Fig 6J). This indicates that: i) IL-33 directly activates nociceptors, ii) that nociceptors from mice with endometriosis are more prone to respond to IL-33, and iii) that the degree of IL-33-induced activation is higher in nociceptors from mice with endometriosis. For better visualization, traces of the IL-33-responsive DRG neurons are also shown in Fig S3. Altogether, our data further supports the notion that IL-33 mediates pain through direct activation of ST2^+^ nociceptors and unveils a previously unknown mechanisms for pain generation during endometriosis.

**Figure 6.**
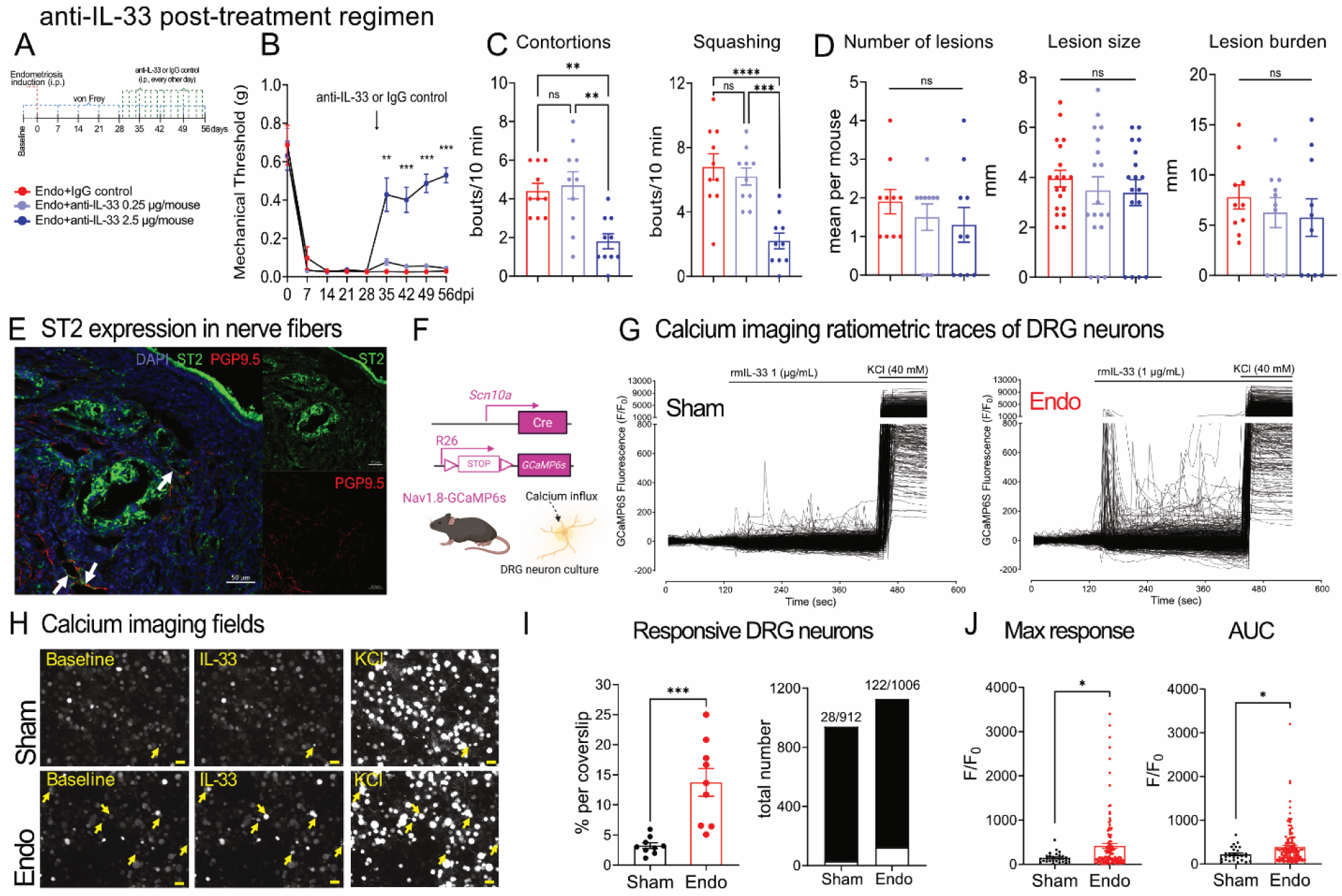
IL-33/ST2 signaling drives endometriosis-associated pain. **(A)** Schematic displaying the disease intervention protocol for the anti-IL-33 antibody. **(B)** Abdominal mechanical hyperalgesia measured before lesion induction (zero) and weekly thereafter using von Frey filaments. Results are presented as mean ± SEM of mechanical threshold, n = 10 mice per group (two-way repeated-measures ANOVA followed by Tukey’s post hoc, ***p<0.001; **p<0.01). **(C)** Spontaneous pain measured by abdominal squashing and abdominal contortion. **(D)** Number of visible lesions calculated as sum of total lesions per mouse while lesion size was measured in the lesions as the average of perpendicular diameters. Lesion burden was calculated as the sum of all lesion size in a given mouse. Results are expressed as mean ± SEM, n = 10 (one-way ANOVA followed by Tukey’s post hoc, ****p< 0.0001; **p<0.01; *p<0.05; n.s. = non-significant). **(E)** Immunofluorescence staining of endometriosis displaying ST2 (green staining, top inset) and PGP9.5 (red staining, bottom inset). Arrows point to ST2^+^PGP9.5^+^ nerve fibers. Scale bar 50 µm. **(F)** breeding strategy to generate Nav1.8-GCaMP6 mice. **(G)** Calcium imaging recordings of cultured DRG neurons from sham (left traces of GCaMP6S fluorescence) and endometriosis lesion-bearing GCaMP6S mice (right traces of GCaMP6S fluorescence). Traces represent the fluorescence intensity of individual neurons over time (10min), relative to their baseline fluorescence (F/F0), which were then normalized to a ratio of 1. Recombinant mouse IL-33 (rmIL-33, 1 µg/mL) and KCl 40mM were applied as indicated. **(H)** Representative images showing neuron’s GCaMP6S fluorescence at baseline (zero-second mark) and following the stimulus with rmIL-33 (1 µg/mL, 120s mark) and KCl (480s mark, activates all neurons). Arrows display neurons responsive to IL-33. Scale 50 µm. **(I)** Grouped data showing the percentage (left bar graph) and total number (right bar graph) of IL-33-responsive DRG neurons, with the white bar representing the number of IL-33-responsive DRG neurons. **(J)** Grouped data showing the overall magnitude of IL-33-induced calcium entry, measured as the peak of GCaMP6S fluorescence (left bar graph), and area under the curve of (AUC, right bar graph). Results were obtained from n = 3 sham and n = 3 endometriosis lesion-bearing GCaMP6S mice; which generated 9 coverslips from each mice type (Mann-Whitney test and unpaired t-test for percentage of DRG neurons, ***p< 0.001; *p<0.05).

## DISCUSSION

To understand the endogenous role of IL-33 and its regulation during endometriosis, we first used unbiased techniques to discover that nociceptor-derived CGRP induces the expression and release of IL-33 by macrophages. We then employed a series of genetic and pharmacological manipulations to prevent IL-33 signaling. As IL-33 is the sole agonist for ST2 (*12, 34, 45*), we specifically determined the extent to which IL-33/ST2 signaling is required for lesion formation by using mice lacking *St2* as tissue donors during endometriosis induction. We further confirmed that data by using treatment with monoclonal antibodies against IL-33 or ST2. To then understand how neuroimmune communication plays a role in IL-33 production, we applied a series of chemical and genetic approaches to ablate nociceptors and macrophages. Altogether, we unveil a hitherto unknown mechanism for neuroimmune-driven production of IL-33 as well as its potential dual role during endometriosis. Using scRNAseq and bulk RNAseq of mouse and human lesions, we identified a nociceptor to PEM pathway that drives IL-33 production. CGRP-driven production of IL-33 by PEMs is key to lesion growth and pain. This is in agreement with previous works showing a CGRP-mediated production of IL-33 by mesenchymal stromal cells (*46*) and innate lymphoid cells (ILC2s) (*47*). Interestingly, these IL-33^+^ mesenchymal stromal cells (*48*) and ILC2s (*49*) seem to be in close proximity to neurons, indicating that neuronal-derived molecules might be one of the regulators of IL-33 in these cells, as well.

In humans, we found that IL-33 correlates with lesion fibrosis and gland numbers, indicating that IL-33 might be associated with both epithelial proliferation and fibrosis in humans. The latter observation is consistent with the ability of IL-33 to drive fibrosis in different organs (33). Furthermore, we also determined that *IL33* transcripts in macrophages are associated with endometriosis risk in humans, as well as the different types of endometriosis lesions such as superficial, deep, and endometriomas as determined by scDRS. In these data, *IL33* was also expressed in several other immune cell clusters such as dendritic cells, plasma cells, effector CD8 T-cells, suggesting that these cells could be also associated with endometriosis risk as well as the different types of lesions in patients. It remains to be determined whether ablating IL-33 from these cell types will play a role in endometriosis. A limitation of our scDRS approach is that the single cell data sets only capture the top 10-20% of genes based on absolute expression, and IL-33 expression, as is typical for cytokines, was only captured in a subset of cells. The observations made here, therefore, need to be taken with caution until further validation. Nevertheless, our combined human (IHC, scDRS) and pre-clinical data (behavior studies, cell proliferation, bulk RNAseq, calcium imaging assays, etc) demonstrate that IL-33 release by PEMs drives lesion formation and pain. We also provided evidence that suggest a dual role for IL-33 in endometriosis. IL-33/ST2 is required for lesion formation as our data demonstrates that uterine horn of mice lacking *St2* generate fewer and smaller lesions. Furthermore, IL-33 induced a direct and potent activation of ST2^+^ nociceptors from endometriosis lesion-bearing mice. Therefore, targeting IL-33/ST2 signaling might be an effective way to treat endometriosis pain.

Our data indicate that IL-33 plays multiple roles in endometriosis pathophysiology. During lesion induction, we found that IL-33 is rapidly released into the peritoneal cavity (peaks at 3 dpi) and then returns to baseline levels. This is consistent with studies showing that in addition to being constitutively produced and released, IL-33 can also be stored in the nucleus. Therefore, after tissue damage IL-33 can be released as an alarmin to initiate immune response (*50, 51*), a process that likely occurs during retrograde menstruation. *In vitro*, we also show that IL-33 induces epithelial cell proliferation, corroborating studies showing IL-33 induces epithelial cell proliferation in skin cancer, colon cancer (*52*), during intestinal injury (*52*), and in nasal mucosal inflammation (*53*). Collectively, our data strongly support a role for IL-33/ST2 signaling in endometriosis lesion formation.

In contrast, anti-IL-33 treatment of established lesions did not significantly reduce lesion size but substantially reduced pain. This points to an ongoing role for IL-33 in pain generation once lesions are fully formed. In agreement, we detected IL-33 levels in the lesions in time points where IL-33 levels in the peritoneal cavity wash had returned to baseline (e.g., 14 dpi). Even though ST2 staining is most prominent in the epithelium of lesions, we also found the presence of ST2^+^PGP9.5^+^ nerve fibers, indicating the presence of IL-33-responding nerve fibers in the lesions. To support those observations, we found that DRG neurons of mice with endometriosis had an increased response to IL-33 when compared to sham mice. This further indicates that IL-33 directly activates nociceptors to trigger pain during endometriosis. Our data confirms data from the literature showing that ST2 is expressed by TRPV1^+^ sensory neurons (*22*) and that it exacerbates disease burden via nociceptor activation (*20–22*). Specifically for pain, our findings corroborates data showing that local injection of IL-33 is sufficient to trigger pain (*16–18, 54*) while injection of suboptimal doses of IL-33 can also potentiate different stimuli to induce pain (*16*). Altogether, we suggest a dual role for IL-33/ST2 signaling during endometriosis. Initially, it is a major contributor to lesion formation but once lesions are fully formed an additional role is added in which throughout lesion lifetime IL-33/ST2 signaling continues to drive pain.

In summary, we identified a neuroimmune-driven production of IL-33 production by PEMs that is key for lesion growth and pain during endometriosis in mice. We also found that IL-33 expression is associated with endometriosis risk in humans. Further, we provide evidence that IL-33 plays dual roles in endometriosis, contributing to both lesion formation as well as ongoing pain. Therefore, targeting IL-33/ST2 signaling might be an effective non-hormonal and non-opioid approach to treating endometriosis.

## METHODS

### Animals

Animal handling and care was conducted in accordance to the United States National Institutes of Health Guide for the Care and Use of Laboratory Animals, and Australian National Health and Medical Research Council (NHMRC), and were in accordance with the laws of the United States and regulations of the Department of Agriculture. All experiments and procedures were approved by the Institutional Animal Care and Use Committee (IACUC) at South Australian Health and Medical Research Institute (protocol number SAM-23-040) for calcium imaging experiments), at Londrina State University (protocols numbers 2669.2019.54 and 047.2023 for ST2-KO experiments), and at Boston Children’s Hospital (protocols numbers 19-12-4054R and 00001816 for all remaining experiments). Healthy C57BL/6 mice (Stock #000664), B6.129-Trpv1^tm1(cre)Bbm^/J (TRPV1cre, stock #017769), B6.129P2-Gt(ROSA)26^Sortm1(DTA)Lky^/J (ROSA-DTA, stock #009669), C57BL/6-Tg(Csf1r-cre)1Mnz/J (Csf1r-cre, stock #029206), and B6;129S6-Gt(ROSA)26Sortm96(CAG-GCaMP6s)Hze/J (GCaMP6s, stock #024106) were purchased from Jackson Laboratories. C57BL/6N-*Ramp1^tm1c(EUCOMM)Wtsi^*/H (RAMP1^fl/fl^) mice were kindly provided by Dr. Isaac Chiu. Csf1r-cre mice were bred with RAMP1^fl/fl^ to generate C57BL/6-Tg(Csf1r-cre)1Mnz/J^+/-^;RAMP1^fl/fl^ that were then bred with RAMP1^fl/fl^ to generated mice with deletion of RAMP1 on macrophages (Csf1r^ΔRamp1^) BALB/c and ST2^-/-^ (ST2-KO, BALB/c background) were received from Ribeirão Preto Medical School (University of São Paulo, Ribeirão Preto, SP, Brazil). B6.129(Cg)-Scn10a^tm2(cre)Jwo/Tjp^J mice (Nav1.8cre, stock #036564) mice were a gift from W. Imlach, Monash University, Australia. Nav1.8cre mice were bred with GCaMP6s mice to generate Nav1.8-GCaMP6s or LM control mice. Female age-matched mice from 8 to 14 weeks of age were used for experiments involving transgenic mouse strains while 8-week-old mice were used for the experiments performed in C57BL6/J mice.

Block randomization was used to randomize subjects into groups resulting in an equal sample size at all time-points. TRPV1 nociceptor ablation was performed as previously described (*55*). For subcutaneous RTX (cat #R400, Alomone), escalating doses of RTX (30, 70, 100 µg/kg on consecutive days) or vehicle (2% DMSO, 0.15% Tween-80 in sterile PBS) were used. Mice were under isoflurane anesthesia. For chemical depletion of macrophages, clodronate liposomes or vehicle control liposomes (Encapsula Nanaoscience) at 0.0625 mg/mouse (*56*) were injected intraperitoneally 24h before endometriosis induction. Mice were randomly assigned and housed in standard clear plastic cages with no more than 5 mice per cage in a 12:12h light/dark cycle with *ad libitum* access to water and food. Behavioral testing was performed between 9 a.m. and 5 p.m. by an investigator who was blind to treatment group in a room maintained at a temperature of 21±1°C. Blinding was maintained until all data analysis decisions were final. All efforts were made to minimize the number of animals used and their suffering. Euthanasia was performed by controlled CO2 inhalation.

### Human sample collection

Biopsies of endometriotic lesions were obtained from patients with a laparoscopic diagnosis of endometriosis at St. Marien Hospital, Amberg, and St. Hedwig Clinic, Regensburg, Germany. The samples were collected during days 3-5 of the menstrual cycle, corresponding to the proliferative phase. This study was approved by the Ethics Committee of the University of Regensburg, Germany (IRB approval numbers 21–2427–101, 23-3211-101, 22-2862-104). Written informed consent was obtained from all patients. For more information, please refer to the “human sample collection and demographic” section in the supplementary materials.

### Statistical analysis

Results are presented as mean ± SEM and followed ARRIVE 2.0 guidelines (*57*). Data were analyzed using the GraphPad Prism version 10.5 software. Two-way repeated measure analysis of variance (ANOVA), followed by Tukey’s *post hoc*, was used to analyze data from experiments with multiple time points (von Frey). One-way ANOVA followed by Tukey’s *post hoc* was used to analyze data from experiments with a single time point. To explore potential associations among IL-33 expression, glandular proliferative status, extent of fibrosis, duration of symptoms, revised American Society for Reproductive Medicine (rASRM) score, worst pain score, age at surgery, and age at menarche, a Spearman correlation analysis was conducted. Comparison between two groups was conducted using two-tailed Student’s t-test. Results are presented as Spearman’s correlation coefficients (ρ). For all analysis, differences were considered significant when p < 0.05.

## ACKNOWLEDGMENTS

We thank Neurodevelopmental Behavioral Core (NBC) of the Boston Children’s Hospital. We also thank Londrina State University’s core facility CMLP-UEL (Central Multiusuário de Laboratórios de Pesquisa da Universidade Estadual de Londrina) for the use of instruments free of charge. F.S.R.-O. thanks the National Council for Scientific and Technological Development (CNPq, Brazil) for the 12-month scholarship to develop her split PhD fellowship (Doutorado Sanduiche, SWE; award number 203112/2020-2). M.D.V.S. thanks the Coordination for the Improvement of Higher Education Personnel (CAPES, Brazil) for the 9-month scholarship to develop his split PhD Fellowship (Doutorado Sanduiche PDSE). F.S.R.-O. and M.D.V.S. thank PhD scholarship from CAPES (finance code 001). W.A.V.J. acknowledges the CNPq Productivity fellowship (award number 309633/2021). We also thank Dr. Ray Anchan for insightful discussions during experimental planning and the preparation of this manuscript. Author contributions: **Conceptualization:** V.F. and M.S.R. **Investigation:** V.F., F.S.R.-O., S.O., E.G., M.V.B., T.A., M.V., C.K., M.E.S., M.D.V.S, O.K.H., J.C. **Formal analysis:** V.F., F.S.R.-O., S.O., E.G., M.V.B., T.A., M.V., C.K., M.E.S., M.D.V.S, O.K.H., J.C. **Data curation:** V.F., S.O., E.G., M.V.B., T.A., M.V., C.K., M.E.S., S.H., J.C, K.L., and M.S.R. **Methodology:** V.F., E.G., M.V.B., T.A., M.V., C.K., M.E.S, W.A.V.J., S.H, J.C., K.L., and M.S.R. **Resources:** V.F., M.V.B., T.A., M.V., C.K., M.E.S, W.A.V.J., S.H., J.C, K.L., and M.S.R. **Project administration:** V.F. **Supervision:** V.F., M.S.R. **Visualization:** V.F., W.A.V.J., and M.S.R. **Writing (original draft):** V.F. **Writing (reviewing and editing):** all authors. All authors read and approved the final version of the manuscript. **Funding acquisition:** V.F., S.M.B., J.C., K.L. and M.S.R. **Data availability statement:** The data that support the findings of this study are available from the corresponding author upon reasonable request. **Conflict of interest:** V.F. and M.S.R. are inventors on patent PCT/US2023/079238, which covers some of the material in this paper. No other authors have conflict of interest to declare.

---

**Figure S1.**
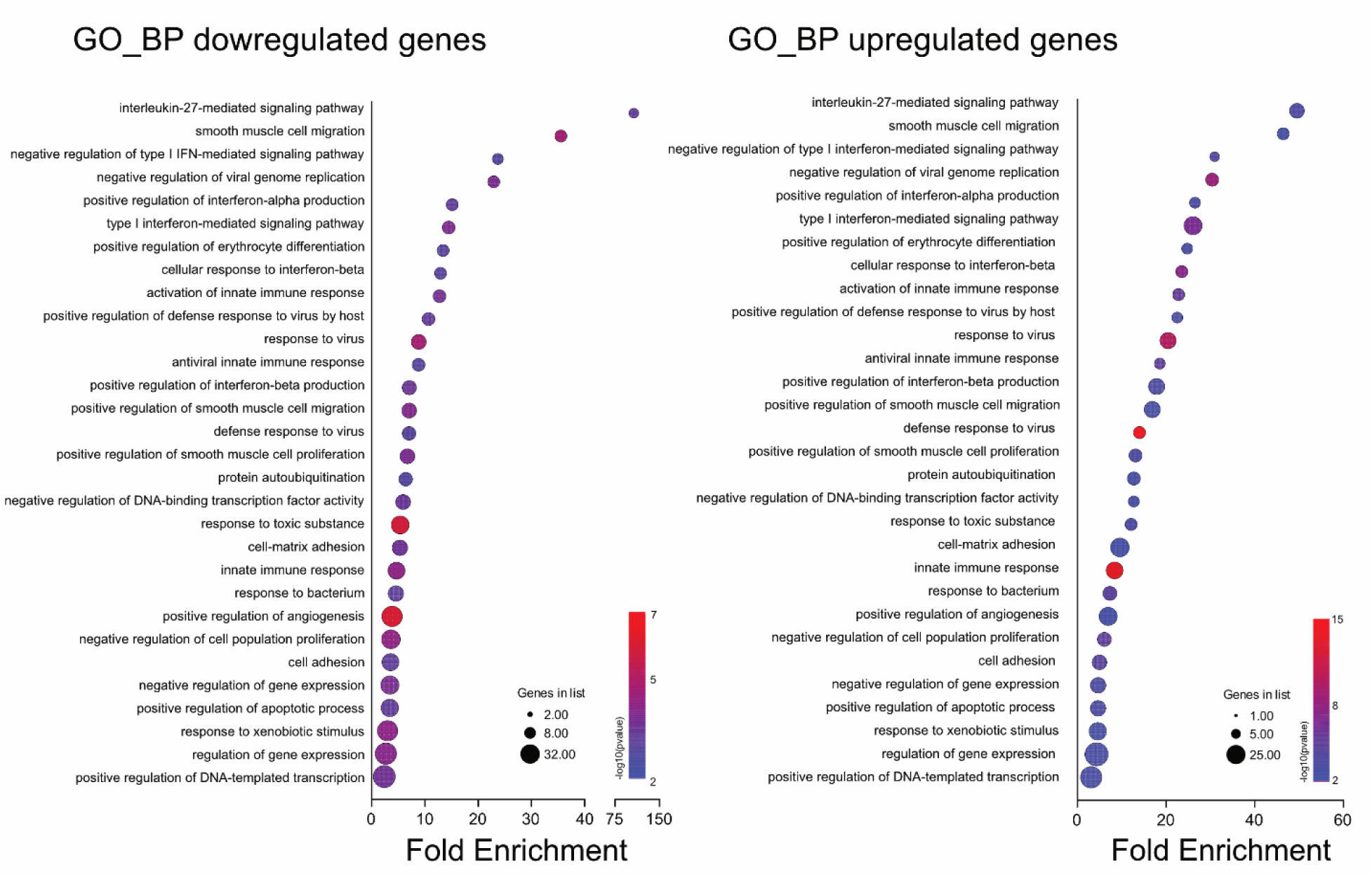
GO enrichment analysis of IL-33-stimulated endo-epi cells. GO enrichment analysis of the downregulated (left) and upregulated (right) genes in endo-epi cells upon IL-33 stimulation (n = 2 for each condition).

**Figure S2.**
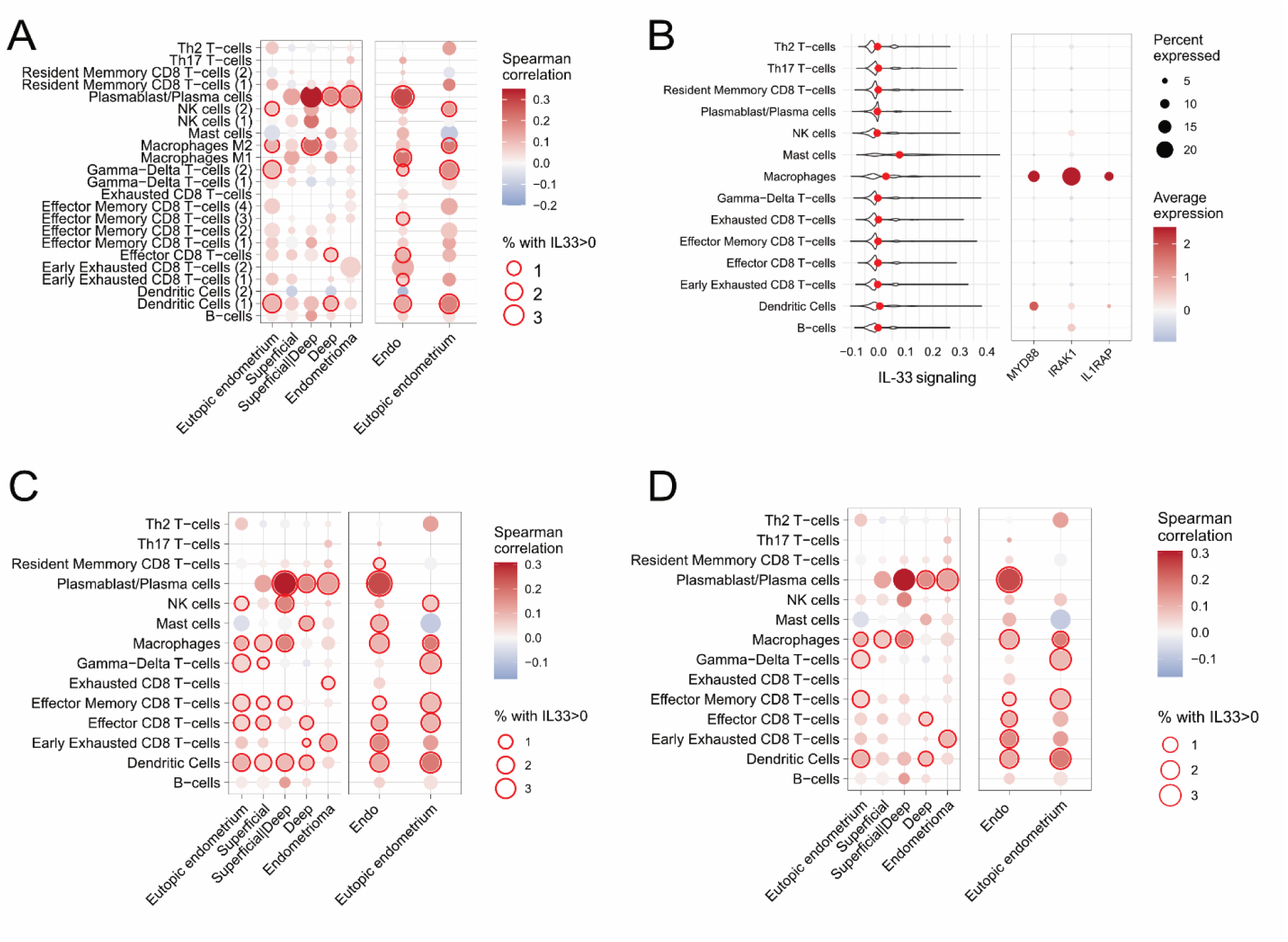
Correlation between IL-33 expression and endometriosis risk. **(A)** Expression of IL-33 in the immune cell clusters with endometriosis lesion subtype risk. Red border highlights significant raw score for IL-33 and the different types of endometriosis lesions and eutopic endometrium. **(B)** Transcriptional signature of IL-33-mediated signaling per cell type. Red dots mark the mean of the distribution. Right table displays selected IL-33/ST2 downstream signaling pathways markers. **(C and D)** Expression of IL-33 in the immune cell clusters with disease risk (aggregating lesion subtypes as “endo”). Red border highlights significant raw and z-score for IL-33 and the presence of endometriosis lesions (endo). Correlation was not calculated for clusters below 50 cells and significance was considered when p-adjusted<0.05 (A and D) or p<0.05 (C).

**Figure S3.**
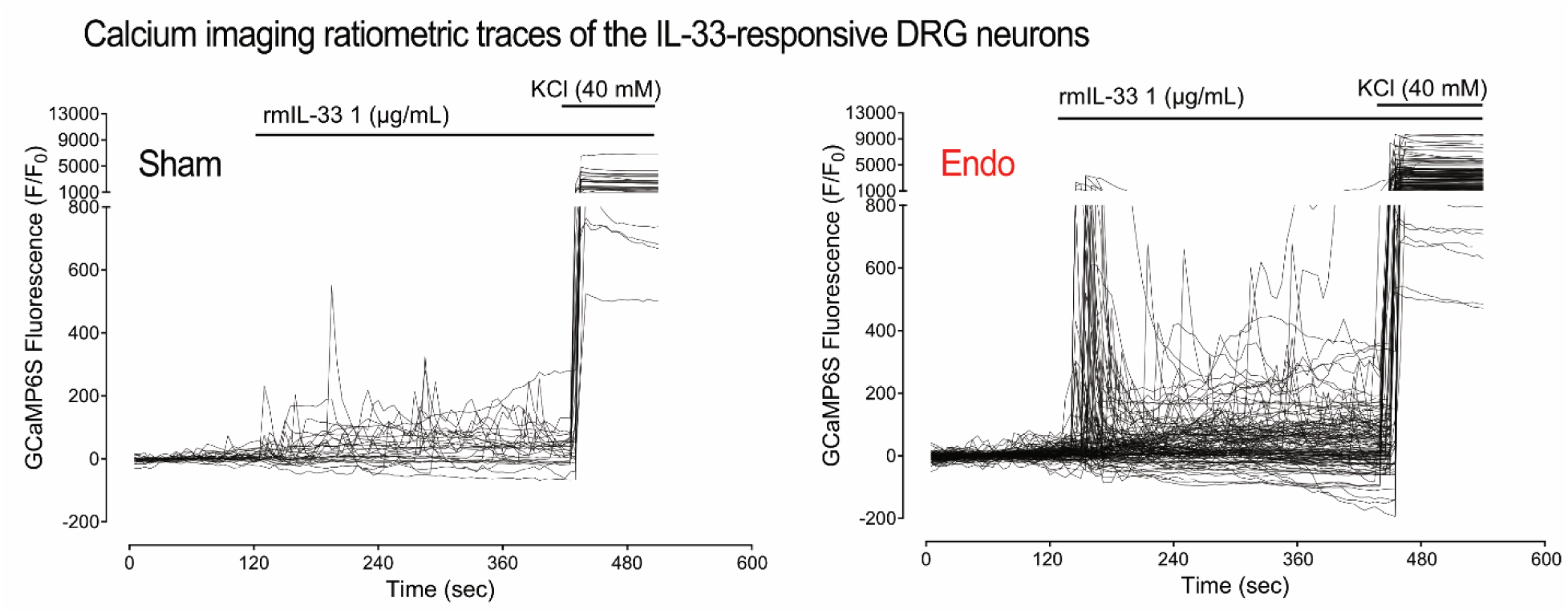
IL-33 directly activates nociceptors during endometriosis. Calcium imaging recordings of cultured DRG neurons from sham (left) and endometriosis lesion-bearing (right) GCaMP6S mice. Fluorescence traces showing individual IL-33-responsive DRG neurons only (28 in sham mice vs 122 in endometriosis mice). Recombinant mouse IL-33 (rmIL-33, 1 µg/mL) and KCl 40mM were applied as indicated. Results are expressed as fluorescence intensity of individual neurons over time (10 min), relative to their baseline fluorescence (F/F_0_), which were then normalized to a ratio of 1. Results were obtained from n = 3 sham and n=3 endometriosis lesion-bearing GCaMP6S mic; which generated 9 coverslips from each mice type.

