## Supplementary material for "Pro-endometriosis macrophage release of IL-33 is key for endometriosis pain and lesion formation": Supp material

#### Human sample collection and demographics

Biopsies of endometriotic lesions were obtained from patients with a laparoscopic diagnosis of endometriosis at St. Marien Hospital, Amberg, and St. Hedwig Clinic, Regensburg, Germany. The samples were collected during days 3-5 of the menstrual cycle, corresponding to the proliferative phase. This study was approved by the Ethics Committee of the University of Regensburg, Germany (IRB approval numbers 21–2427–101, 23-3211-101, 22-2862-104). Written informed consent was obtained from all patients. The diagnostic of endometriosis patients was performed as described in our previous study (*58*), following the revised American Society for Reproductive Medicine classification criteria (rASRM) that stages the extent and severity of endometriosis based on the size, location, and depth of lesions, as well as the presence and extent of adhesions. All participants were premenopausal, presented with typical endometriosis-related symptoms (e.g., pain and/or sterility), and underwent laparoscopic surgery for the removal of endometriotic lesions. Data processing and analyses were conducted in accordance with the Declaration of Helsinki and the General Data Protection Regulation of the European Union. Patient demographics and pain score are presented in Table S1.

#### Single cell Disease Relevance Score (scDRS)

Inherited risk data was integrated with human single cell transcriptomics with the single cell disease relevance score software (*35*). Data comes from studies GCST90205183 (*30*) and GSE213216 (*32*). Spearman correlation between the score and *IL33* expression was estimated per cell type and disease state for all cell groups with more than 50 cells. False discovery rate was used for multiple test correction, using 0.05 as significance threshold. IL-33 signaling was evaluated by aggregating the expression of markers in the gene ontology process GO:0038172 with function AddModuleScore from the Seurat R package (*35*).

#### Induction of endometriosis in the mouse model

After at least one week of acclimatization, donor mice received a subcutaneous injection of 3 μg/mouse estradiol benzoate (30 μg/mL in sesame oil) to stimulate the growth of the endometrium as previously described (*59*).

The uteri of donor mice were dissected into a Petri dish containing sterile Hank's Balanced Salt Solution (HBSS) and minced with scissors and scalpel one at the time, ensuring that the maximal diameter of each fragment was consistently smaller than 1 millimeter (mm). Each dissociated uterine horn was then injected intraperitoneally into a recipient mouse in 500 μL of sterile HBSS, meaning that one uterus of a donor mouse was used for endometriosis induction in every two recipient mice. Sham mice received an intraperitoneal injection of 500 μL of sterile HBSS.

For experiments involving calcium imaging using Nav1.8-GCaMP6S mice, we used our previously established mouse model of surgically induced endometriosis (*60*). Briefly, Nav1.8-GCaMP6S mice female mice were ovariectomized to deplete endogenous steroid production and administered with an intraperitoneal (i.p.) injection of 100 µg/kg estradiol benzoate to maintain steady levels of circulating estrogen and minimize any difference related to the stage of the estrous cycle. Seven days later, mice were anesthetized, and the uterine horns were exposed, excised and transferred into a dissecting dish containing ice-cold sterile PBS supplemented with penicillin (100 U/ mL) and streptomycin (100 µg/mL, Sigma-Aldrich). Uterine horns were longitudinally opened, with 5 × 3 mm fragments of uterine horn tissue removed using a 3 mm biopsy punch (Kai Medical, cat #KAI00010). Three fragments were sutured back on alternate mesenteric cascade arteries that supply the small intestine, and the remaining two pieces were sutured on either side of the mouse's uterus dome. All mice were given an i.p. injection of 100 µg/kg estradiol benzoate immediately after surgery, and once a week for up to 10 weeks, as per (*60*). Sham/control mice were generated using the same procedure, but fat (instead of uterine horn tissue) was sutured in the same locations. We have previously characterized this mouse model of endometriosis and showed that they developed endometriotic lesions, neuroangiogenesis and an enhanced inflammatory environment within the peritoneal cavity. These mice also displayed enhanced sensitivity to visceral and cutaneous pain and altered bladder function (*60*).

**Mouse sample immunofluorescence**

Lesions or dorsal root ganglion (DRG) neurons samples were dissected at 28 days post induction (dpi) for immunofluorescence. Lesions or paired thoracic and lumbosacral (T10-L3 plus L6-S1) DRG were dissected and maintained in 4% paraformaldehyde (PFA) for 24h and then transferred to 30% sucrose for 72h, then to optimum cutting temperature reagent (Tissue-Plus O.C.T., Thermo Fisher Scientific). Embedded samples were cut in 16 µm sections while peritoneal cavity wash cells were mounted on slides with Fluoromount-G compound (cat #0100-01, SouthernBiotech). Primary antibodies used in this study are as follow: anti-PGP9.5 (1:50, cat #ab8189, Abcam), anti-ST2 (1:100, cat #Ab25877, Abcam); anti-IL-33 (10 µg/mL, cat #AF3626, R&D Systems); anti-F4/80 (1:500, cat #14-4801-82, Life Technologies); Ki-67 (1:50, cat #Ab279653, Abcam). All primary antibodies and DAPI (2 µg/ml; 1:500, cat #62248, Life Technologies) were incubated overnight at 4 °C (2.5% goat serum in 0.3% PBS-Tween-PBST). Secondary antibodies used were: goat anti-rabbit Alexa Fluor 488 (1:1000, cat #A-11029, Life Technologies); goat anti-mouse Alexa Fluor 647 (1:500, cat #A-21236, Life Technologies); goat anti-mouse 594 (1:200, cat #DI-2594, Vector); donkey anti-goat Alexa Fluor 488 (1:1000, cat A32814, ThermoFisher); donkey anti-rat Alexa Fluor 647 (1:500, cat #A78947, ThermoFisher) and incubated for 50 min at room temperature (2.5% goat serum in 0.3% PBST). Stained slides were then washed five times with 0.3% PBST and mounted in Fluoromount-G compound (cat #0100-01, SouthernBiotech). Images were taken and processed on Zeiss LSM 880 laser scanning microscope using 20x objective (Carl Zeiss Microscopy). Fluorescence intensity was measured using the measure function on ImageJ (NIH). The level of co-localization for Ki67-ST2 and F4/80-IL-33 experiments was performed using the co-localization feature in Zen Blue software (Carl Zeiss Microscopy).

**Human sample immunohistochemistry and staining.**

Paraffin-embedded human endometrial ectopic lesions (5 µm-thick sections) were stained with primary antibodies against Ki-67 (proliferation) or IL-33. Briefly, sections were deparaffinized, rehydrated, and antigens were retrieved using 0.01 M sodium citrate buffer (pH 6.0, cat #C9999, Sigma). Endogenous peroxidase activity was blocked with 3% $H_2O_2$ (cat #107209.0250, Sigma) for 5 minutes at room temperature. Sections were incubated overnight at 4 °C with primary antibodies against IL-33 (1:200, cat #12372-1-AP, Proteintech) or Ki67 (1:250, cat #ab16667, Abcam). Negative controls were treated with antibody diluent (cat #S3022, DAKO) instead of the primary antibody. After washing, a goat anti-rabbit IgG secondary antibody (1:200, cat #65-6140; Invitrogen) and streptavidin (cat #ab64269, Abcam) were applied. Staining was visualized using diaminobenzidine (DAB, cat #ab64238, Abcam), and sections were counterstained with hematoxylin. Images were acquired using a slide scanner (cat #P1000, 3DHistech) and CaseViewer (v2.3) software. IL-33 expression was quantified by measuring the integrated optical density (IOD) in three representative visual fields (VFs) surrounding glands and three within fibrotic regions using ImageJ (v1.53k). Ki67$^{+}$ epithelial cells within the glands were manually counted and quantified as a percentage of the total epithelial cell population using CaseViewer (v2.3). When possible, Ki67 staining was performed on consecutive tissue sections from the same samples used for IL-33 analysis. Although all glands were IL-33$^{+}$ and it was possible to match the IOD of IL-33 in glandular areas with the percentage of Ki-67$^{+}$ cells from the same tissue, this does not represent exact co-localization of the two markers. Tissue sections were stained with Masson-Goldner trichrome to detect collagen deposition characteristic of fibrosis. Sequential incubation in hematoxylin, ponceau, phosphotungstic acid, and light green solutions was performed, with 1% glacial acetic acid enhancing staining between steps. Collagen-rich connective tissue appeared green, cytoplasm red, and erythrocytes orange. All analyses were conducted by an investigator blinded to the groups.

**DRG sensory neurons cell culture for calcium imaging recordings**

Nav1.8-GCaMP6s mice with fully developed endometriosis (8-10 weeks after model generation) or sham mice, were humanely euthanized via $CO_2$ inhalation, and thoracolumbar (T10-L3) and lumbosacral (L5-S1) dorsal root ganglia (DRG) were removed. DRGs were enzymatically digested in Hanks' balanced salt solution (HBSS; pH 7.4; Life Technologies, cat #14170161) containing 4 mg/mL collagenase II (GIBCO, ThermoFisher Scientific, cat #17101015) and 4 mg/mL dispase (GIBCO, ThermoFisher Scientific, cat #17105041) at 37°C for 30 min. A subsequent incubation (10 min at 37°C) with HBSS containing 4 mg/mL collagenase was further applied. DRGs were washed two times with HBSS and then triturated through a fire-polished Pasteur pipette until a single-cell suspension was achieved, as previously described (*44, 61*). The cell suspension was centrifuged at 50 g for 1 min, and the cell pellet was resuspended in DMEM (25 mM glucose, 1 mM pyruvate, Gibco) containing 10% FCS (Invitrogen), 1x GlutaMAX (Gibco), 1x MEM non-essential amino acids (Gibco), 1x penicillin/streptomycin (Invitrogen), and 10 ng/mL NGF (Merck). DRG neurons were spot-plated on coverslips coated with poly-D-lysine (Merck, 100 µg/mL) and laminin (Merck, 18 µg/mL) and maintained at 37 °C in 5% CO2 for 20-28 hours, for calcium imaging assays, as previously (*44, 61*).

**Calcium imaging on cultured DRG sensory neurons**

After ~20 hours of plating, coverslips contained cultured DRGs were transferred to a recording chamber filled with Ringer's solution (NaCl 140 mM, KCl 5 mM, $CaCl_2$ 1.25 mM, $MgCl_2$ 1mM, glucose 10 mM, HEPES 10 mM, pH 7.4) at room temperature (~23°C). Intracellular calcium changes were measured by recording changes in GCaMP6S florescence at room temperature (23°C), as previously described (*44*). Briefly, GCaMP6S was excited at 480 nm, and emission was measured at 530 nm using an Olympus IX71 microscope in conjunction with a Sutter Lambda 10-3 wavelength switcher, and the Chroma filter set no. 49011 (ET480/40x (Ex), T510lpxrxt (BS), ET535/50m (Em)). Fluorescence images were obtained every 5 seconds, using a X10 objective, with a monochrome CCD camera (Retiga ELECTRO). Images were taken at baseline and following 5 min incubation with recombinant mouse IL-33 (1µg/mL; Bio-Techne, Cat no. 3626-ML-010/CF). After the IL-33 incubation period, neuron's viability and maximum levels of intracellular calcium were verified by incubation with 40 mM KCl. Fluorescence traces of cell bodies were recorded and analyzed offline using MetaFluor Imaging Software (Molecular Devices, CA). Regions of Interest (ROIs) were manually drawn around the cell bodies of neurons. The fluorescence intensity within each cell over time was normalized relative to the baseline fluorescence ($F/F_0$) and then normalized to a ratio of 1. Grouped data is presented as mean ± SEM, with normality of data assessed using the Shapiro-Wilk test prior to all statistical analysis. Grouped data was analyzed using Unpaired T-test and a Mann-Whitney test for nonparametric data. For analysis, neurons responding to IL-33 were identified as neurons with an increase in GCaMP6s fluorescence of more than 10% from baseline fluorescence. The degree of the response to IL-33 was quantified using Prism software (v.10.5,0, GraphPad Software, San Diego, CA, USA), by calculating the maximum amplitude (peak of GCaMP6s fluorescence) and the area under the curve (AUC) of the response to IL-33. Differences were considered statistically significant at $P < 0.05$.

**Behavioral testing**

For mechanical hyperalgesia test, mice were allowed to habituate to the apparatus for at least 2h and during three consecutive days before the beginning of measurements as previously described (*59*). The mechanical threshold was determined by the up and down method starting with 0.4g filament and calculated using the open source software Up-Down Reader (*62*).

For spontaneous abdominal pain measurements, stretching the abdomen (abdominal contortions) and squashing of the lower abdomen against the floor were quantified as previously described (*59*). Briefly, for abdominal contortions, mice were placed in individual chambers in a temperature-controlled (29 °C) glass plate and the number of abdominal contortions was quantified for 10 min. Positive responses consist of a contraction of the abdominal muscle together with stretching of hind limbs. For abdominal squashing, the number of times the mice pressed the lower abdominal region against the floor in 10 minutes was quantified (*59*). Doses for the FDA-approved drugs used here were based on our previous paper (*11*). Doses for anti-IL-

33 antibody (cat #AF3626, R&D Systems) were based on previous works (63, 64). Investigators were blinded to the treatments at all times.

**IL-33 ELISA**

Lesions or uterine horn were dissected at 14 dpi (Fig 5A), 28 dpi (Fig 4A and C; Fig 5D), 42 dpi (Fig 4B, rimegepant cohort), or 56 dpi (Fig 4B, other four drugs) into lysis buffer (PBS, 1% Triton X-100 and protease inhibitors) and then homogenized and centrifuged (10,000 g × 5 min, 4 °C). PerC wash was collected from sham and lesion-bearing mice at 3, 7, 14, 21, and 28 dpi using 2 mL of buffer containing phosphate-buffered saline (PBS), and 2.5 mM ethylenediaminetetraacetic acid (EDTA). To generate the uterine horn of the donor mice, 3 µg/mouse estradiol benzoate (30 µg/mL in sesame oil) to stimulate the growth of the endometrium as previously described (*59*). The ELISA kit was purchased from ThermoFisher (cat #50-112-5200).

**Lesion size and lesion number quantification**

At the specified timepoints, lesions were carefully dissected and measured using a caliper. Lesion size is expressed in millimeters (mm) calculated as the average of two measurements (width and height) since this measure is approximately normally distributed. The number of lesions per mouse was determined by a simple count of the visible lesions (*59*).

**Cell culture**

Peritoneal cavity washes were collected into FACS buffer (phosphate-buffered saline [PBS], 0.5% bovine serum albumin (BSA), and 2.5 mM ethylenediaminetetraacetic acid [EDTA]) and cells were centrifuged for 10 min at 300g of naïve C57BL/6 mice. Peritoneal cavity immune cells were seeded in petri dishes in DMEM-F12 media and incubated at 37 °C in 5% $CO_2$ for 60 minutes. Non-adherent cells were discarded, and adherent cells (macrophages) were scraped, centrifuged, and counted. For experiments involving endo-epi cell proliferation, macrophages were plated (100,000 cells/well) onto tissue culture insert and left overnight with 20,000 endo-epi cells (passage 23) plated on the bottom of the well. Stimulation with CGRP (100 nM, cat #RP11095, Genscript) or vehicle was performed for 24 hours, and then endo-epi cells were used to determine cell proliferation assay using a CyQuant™ kit (cat #C7026, Life Technologies) in a BioTek Synergy H1 multimode reader (Agilent). Treatment with rimegepant (100 nM, cat #HY-15498, MedChemExpress LLC) or vehicle was performed 30 minutes before stimulus with CGRP. For IL-33 related experiments, endo-epi cells were stimulated with rmIL-33 (0.002 – 20 ng/mL, cat #3625, R&D Systems) or vehicle for 24h. Treatment with anti-ST2 (0.001 – 10 µg/mL, cat #MAB10041, R&D Systems) or IgG control was performed along with IL-33 stimulus.

**Bulk RNA sequencing and ligand/receptor interaction analysis**

Bulk RNAseq of vehicle- or CGRP-stimulated PEMs was re-analyzed from our previous manuscript (*11*). For the IL-33-stimulated endometrial epithelial cell (endo-epi cells), cells were stimulated with vehicle or recombinant mouse IL-33 (2 ng/mL) for 24 hours. After that, cells (viability >98%) were harvested and RNA was extracted using PureLink™ RNA Mini Kit (cat #12183025, Life Technologies). Sequencing was conducted at Novogene, Inc. Differential gene expression analysis was performed in sequenced datasets using the DESeq2 via Partek™ Flow™ software (version 12.8.0). Genes were considered differentially expressed when the adjusted p-value was <0.05. The list of differentially expressed genes was used in pathway enrichment analysis and to create volcano plots using Partek™ Flow™ software.

CellPhoneDB (version 4.1) was used to determine ligand/receptor interaction between CGRP-stimulated macrophages and endo-epi cells (*65*). A manual search of the upregulated genes that can be secreted (e.g. chemokines and cytokines) in CGRP-stimulated macrophages was performed to determine their receptors. Subsequently, we looked for the raw expression (transcript per million as per our bulk RNAseq) of these receptors in the endo-epi cells to create the table in Fig 1C.

**Table S1. Patient demographics.** Worst pain score (WPS) refers to the highest pain intensity reported by the patient during the initial assessment, regardless of whether it was described as dysmenorrhea, dysuria, or dyspareunia, as measured using a visual or numerical rating scale. n.s. = non specified. Mean ± standard deviation (μ ± ς). n = number of patients.

| | μ ± σ | Total (n=13) |
|---|---|---|
| **Age at Surgery (yrs)** | 34.4 ± 4.5 | 13 |
| **Symptom Onset** | | |
| < 6 months | | 2 |
| 1-3 years | | 2 |
| 3-10 years | | 2 |
| > 10 years | | 2 |
| n.s. | | 5 |
| **Endometriosis extent and severity (rASRM stage)** | 2.2 ± 1.0 | |
| 1 | | 4 |
| 2 | | 3 |
| 3 | | 5 |
| 4 | | 1 |
| **Received hormonal therapy** | | |
| yes | | 2 |
| no | | 9 |
| n.s. | | 2 |
| **Tissue origin** | | |
| peritoneum | | 12 |
| ovary | | 1 |
| **Patient ID** | | **Worst pain score** |
| 01 | 7.5 ± 2.1 | 7 |
| 02 | | 8 |
| 03 | | 5 |
| 04 | | 9 |
| 05 | | 3 |
| 06 | | 10 |
| 07 | | 10 |
| 08 | | 9 |
| 09 | | 8 |
| 10 | | 5 |
| 11 | | 7 |
| 12 | | 8 |
| 13 | | 9 |

### Table S2. IL-33 staining correlations

| | | | Age_at_menarche | Age_at_surgery | Worst_Pain_NRS_Scale.0 | Symptom_onset | rASRM | IL33 IOD_around_glands | IL33 IOD_fibrotic_area | IL33 IOD_total_glands | Score_fibrosis | %_Ki67_glands | #_glands |
|---|---|---|---|---|---|---|---|---|---|---|---|---|---|
| Spearman's rho | Age_at_menarche | Correlation Coefficient | -- | | | | | | | | | | |
| | | Sig. (2-tailed) | | | | | | | | | | | |
| | | N | 10 | | | | | | | | | | |
| | Age_at_surgery | Correlation Coefficient | 0.460 | -- | | | | | | | | | |
| | | Sig. (2-tailed) | 0.181 | | | | | | | | | | |
| | | N | 10 | 16 | | | | | | | | | |
| | Worst_Pain_NRS_Scale.0 | Correlation Coefficient | -0.289 | -0.227 | -- | | | | | | | | |
| | | Sig. (2-tailed) | 0.418 | 0.398 | | | | | | | | | |
| | | N | 10 | 16 | 16 | | | | | | | | |
| | Symptom_onset | Correlation Coefficient | -0.223 | 0.176 | 0.006 | -- | | | | | | | |
| | | Sig. (2-tailed) | 0.596 | 0.627 | 0.986 | | | | | | | | |
| | | N | 8 | 10 | 10 | 10 | | | | | | | |
| | rASRM | Correlation Coefficient | 0.243 | -0.084 | -0.331 | 0.016 | -- | | | | | | |
| | | Sig. (2-tailed) | 0.499 | 0.757 | 0.210 | 0.964 | | | | | | | |
| | | N | 10 | 16 | 16 | 10 | 16 | | | | | | |
| | IOD_mean_around_glands | Correlation Coefficient | -0.029 | 0,102 | -0.667* | -0.018 | -0.430 | -- | | | | | |
| | | Sig. (2-tailed) | 0.956 | 0.795 | 0.050 | 0.969 | 0,247 | | | | | | |
| | | N | 6 | 9 | 9 | 7 | 9 | 9 | | | | | |
| | IOD_mean_fibrotic_area | Correlation Coefficient | -0.404 | 0.022 | -0.658* | 0.368 | -0.268 | 0.719* | -- | | | | |
| | | Sig. (2-tailed) | 0.369 | 0.947 | 0.020 | 0.370 | 0.400 | 0.045 | | | | | |
| | | N | 7 | 12 | 12 | 8 | 12 | 8 | 12 | | | | |
| | IOD_mean_totalgl_fi | Correlation Coefficient | -0.235 | -0.122 | -0.747* | -0.165 | -0.469 | 0.929** | 0.850** | -- | | | |
| | | Sig. (2-tailed) | 0.653 | 0,774 | 0.033 | 0.723 | 0.241 | 0.001 | 0.007 | | | | |
| | | N | 6 | 8 | 8 | 7 | 8 | 8 | 8 | 8 | | | |
| | Score_fibrosis | Correlation Coefficient | 0.273 | 0.277 | -0.266 | 0.671 | -0,274 | 0.627 | 0.516 | 0.546 | -- | | |
| | | Sig. (2-tailed) | 0.553 | 0,383 | 0.404 | 0.069 | 0,389 | 0.096 | 0.086 | 0.162 | | | |
| | | N | 7 | 12 | 12 | 8 | 12 | 8 | 12 | 8 | 12 | | |
| | percentage_Ki67_glands | Correlation Coefficient | 0.441 | -0.380 | 0.503 | -0.065 | 0,516 | -0.623 | -0.380 | -0.599 | -0.192 | -- | |
| | | Sig. (2-tailed) | 0.381 | 0.353 | 0.204 | 0.890 | 0.191 | 0.099 | 0.354 | 0.117 | 0.649 | | |
| | | N | 6 | 8 | 8 | 7 | 8 | 8 | 8 | 8 | 8 | 8 | |
| | Number_glands | Correlation Coefficient | -0.418 | -0.012 | -0.207 | 0.519 | -0.225 | 0.313 | 0.818* | 0.458 | 0.538 | -0.188 | -- |
| | | Sig. (2-tailed) | 0.410 | 0.977 | 0.622 | 0.233 | 0.592 | 0.450 | 0.013 | 0.254 | 0.169 | 0.656 | |
| | | N | 6 | 8 | 8 | 7 | 8 | 8 | 8 | 8 | 8 | 8 | 8 |

*. Correlation is significant at the 0.05 level (2-tailed).
**. Correlation is significant at the 0.01 level (2-tailed).

IOD: integrated optical density
NRS: Numeric Rating Scale
IOD_mean / IOD_totalgl_FI: The sum of the average integrated optical density (IOD) values measured across three selected visual fields.
Worst_Pain_NRS_Scale.0: This refers to the patient's self-reported score for worst pain, assessed using the NRS, where 0 indicates "before surgery" assessment.
rASRM: Revised American Society for Reproductive Medicine classification
